# PhaSeDis: A Manually Curated Database of Phase Separation–Disease Associations and Corresponding Small Molecules

**DOI:** 10.1101/2024.05.01.591645

**Authors:** Taoyu Chen, Guoguo Tang, Tianhao Li, Zhining Yanghong, Chao Hou, Zezhou Du, Liwei Ma, Tingting Li

## Abstract

Biomacromolecules form membraneless organelles through liquid–liquid phase separation in order to regulate the efficiency of particular biochemical reactions. Dysregulation of phase separation might result in pathological condensation or sequestration of biomolecules, leading to diseases. Thus, phase separation and phase separating factors may serve as drug targets for disease treatment. Nevertheless, such associations have not yet been integrated into phase separation related databases. Therefore, based on MloDisDB, a database for membraneless organelle factor–disease association previously developed by our lab, we constructed PhaSeDis, the phase separation–disease association database. We increased the number of phase separation entries from 52 to 185, and supplemented the evidence provided by the original article verifying the phase separation nature of the factors. Moreover, we included the information of interacting small molecules with low or high-throughput evidence that might serve as potential drugs for phase separation entries. PhaSeDis strives to offer comprehensive descriptions of each entry, elucidating how phase separating factors induce pathological conditions via phase separation and the mechanisms by which small molecules intervene. We believe that PhaSeDis would be very important in the application of phase separation regulation in treating related diseases. PhaSeDis is available at http://mlodis.phasep.pro.

## Introduction

Phase separation (PS) is a physiochemical phenomenon where the homogeneous solution of macromolecular components separates into distinct phases, with one enriched with certain macromolecules and the other depleted of such molecules. Among many PS scenarios, liquid–liquid phase separation (LLPS) refers to the separation of two distinct liquid phases like oil and water, which was found to exist in a number of biological context [1]. Ever since the first observation of P granule having PS-like droplet properties, researchers in biology and medicine start to pay close attention to PS as it has the potential to unravel the forming and regulating mechanisms of membraneless organelles (MLOs) [2], such as stress granules[3], paraspeckles[4], and nucleoli[5]. Previous studies have also shown that the dysregulation of PS might be one of the vital mechanisms underlying neurodegenerative diseases or other common diseases like cancer [6,7]. Small molecules partitioned into condensates might also regulate PS, rescuing cells from a pathological state [8,9]. Therefore, understanding the role of PS in human cells, particularly in pathological context, is pivotal for the intervention of PS-related diseases and screening potential small molecules.

Researchers have been working hard in their endeavor towards investigation of biological PS [10,11]. Nevertheless, these researches are scattered, as researchers tend to carry out different experiments in verifying the condensates of interest as “PS-related bodies” [12,13]. Moreover, although there are small molecules like 1,6-hexanediol that have been proven to regulate PS [14], researchers of different background make use of a wide range of different small molecules in affecting PS and PS bodies [15−18]. Recent research indicates that small molecules can concentrate in biomolecular condensates of appropriate chemical environment, which can assist in constructing machine learning models to predict small molecule partitioning in PS bodies [19]. Many pieces of evidence have also stated that small molecules might be capable of interfering with the pathological conditions in cells since drug-applied cell lines can witness disturbance or assembly of phase separating bodies [20,21]. Thus, we believe that collecting small molecule interference examples from literatures concerning valid PS-driven condensates should be of great interest to the researchers in the field.

Several databases have been established for the needs of PS researchers, including PhaSepDB [22], LLPSDB [23], PhaSePro [24], and DrLLPS [25]. However, these databases focus on the nature of possibly phase separating molecules, while neglecting the functions and diseases the PS molecule has relations to [26]. With that in mind, we established the first MLO-related disease database MloDisDB in 2020 [27]. Nevertheless, there has been a few limitations with MloDisDB. To begin with, the number of PS entries are limited, with only 52 entries out of the 775 MLO-related entries, which is a great neglect of PS as the possible mechanisms of the formation of mambraneless organelles. Second, the annotations of small molecule interaction do not present, which is of vital significance for researchers who wish to evaluate the PS properties of different factors and screen potential drugs interacting with the PS-determining region of the protein, reversing the abnormal state of the cell. Thus, in this paper we present PhaSeDis, where we supplemented novel data and annotated the *in vivo* experiments evidence and related small molecules. PhaSeDis contains 931 entries, with 185 PS-related entries. We hope the newly designed database would be of great assistance in the PS field.

## Database implementation

### The new database doubles the number of entries related to phase separation

Our previous MloDisDB was released in 2020, which is the first literature-based database of its kind that integrates the relations of MLO-related biomolecules and diseases. Nevertheless, of all the 775 entries upon the first release, only 52 entries are annotated as “phase separation” related biomolecules, while the rest are correlated with certain membraneless organelles. These years have witnessed an increasing concern of PS and related factors, particularly those affiliated with diseases. As researchers put more effort into investigating the PS properties of disease-related biomolecules, we decided that building a novel database concerning PS-disease associations would be pivotal, which leads to the development of PhaSeDis.

In the current version of PhaSeDis, not only did we inherit previous PS-disease association entries from MloDisDB, we also collected all PS-related publications listed on NCBI PubMed by manually examining the search results of the keyword “(phase separation[Title/Abstract]) AND ((disease[Title/Abstract]) OR (cancer[Title/Abstract]) OR (neurodegeneration[Title/Abstract]))”. Considering that MloDisDB collected data up to April 2020, the time range were set as “2020–2022” to collect newer publications. All publications searched by previous keywords and filters were examined manually in full text for extracting comprehensive annotations. We only filter publications with results showing *in vivo* evidence of factors (proteins or RNAs) phase separating into condensates or granules.

As a result, we collected 553 publications from April 2020 (the end point of the MloDisDB) until December 2022. After going over all the literature, we curated 156 novel entries, 133 of them being PS-related entries. That boosts the total number of PS-related entries from 52 to 185, and the total number of entries from 775 to 931. These 185 entries are derived from 123 papers, in which the authors provided direct experiment, indirect experiment or clinical investigation evidences of PS-related factors affecting the corresponding diseases through PS. We follow the same protocol as in MloDisDB [27] in recording the properties of the phase separating factor (type, name, Gene ID, and Uniprot ID, if being a protein), the information of related diseases (classification, name, disease ontology, Medical Subject Headings (MeSH), International Classification of Diseases 10 (ICD-10), Online Mendelian Inheritance in Man (OMIM), etc.), MLO changes or the changes in phase-separating droplets (size, number, assembly, dynamic, etc.) as well as the description in the paper indicating the mechanism of how PS factor leads to the aforementioned disease (**Figure 1**). In a nutshell, we doubled the PS-disease associations entries in PhaSeDis, which reflects the advancement of recent PS researches.

**Figure 1.**
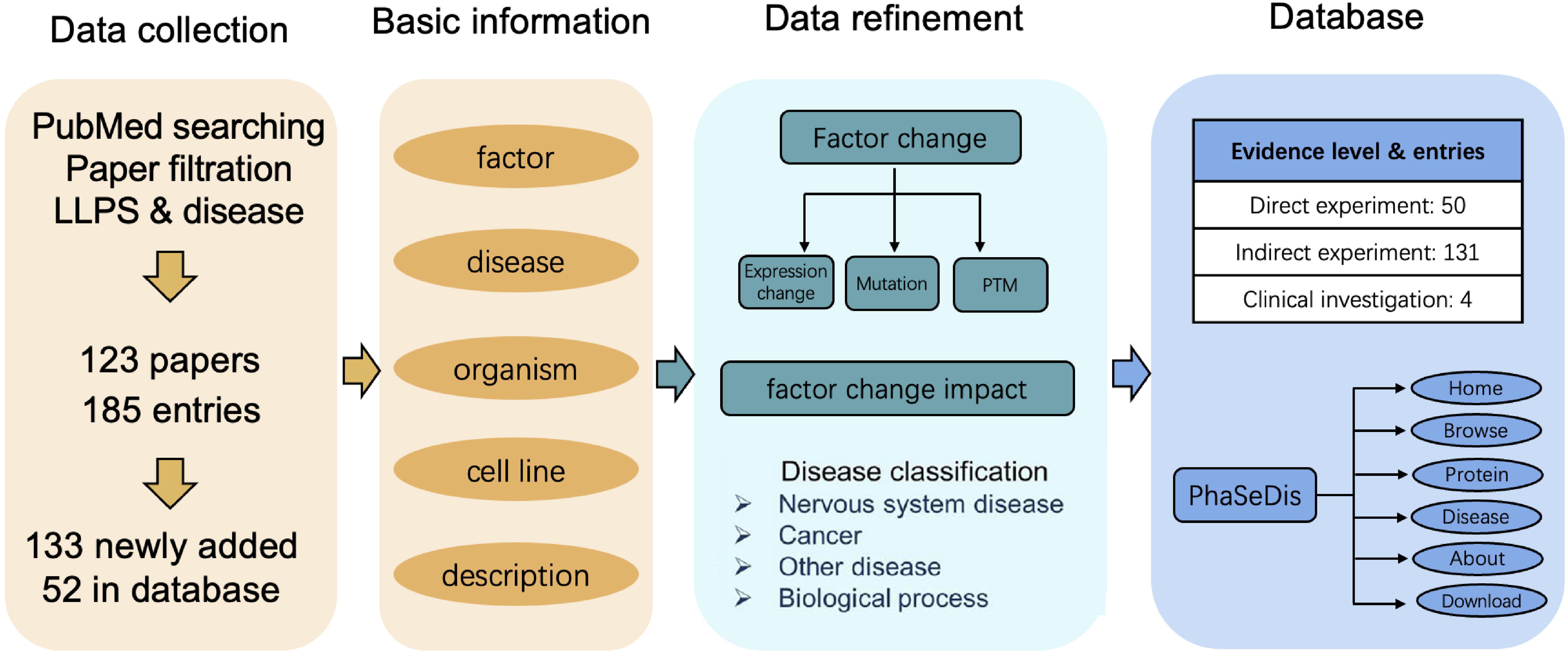
Procedures of PhaSeDis data curation and construction. PhaSeDis data were derived from papers published from year 2020 to 2022 enlisted in PubMed using the search term “(phase separation[Title/Abstract]) AND ((disease[Title/Abstract]) OR (cancer[Title/Abstract]) OR (neurodegeneration[Title/Abstract]))” and previous MloDisDB. We filtered 123 papers and curated 185 entries. These entries were annotated with phase separating factor name, disease, organism, cell line, and a brief description. Then we refined the data with details about expression changes, mutation, and PTM changes along with their impact on pathological conditions, related disease classification, and evidence levels. All aforementioned basic information and data refinements were illustrated in the final database with a web interface. LLPS, liquid–liquid phase separation. PTM, post-translational modification.

Next, we summarized the 185 PS entries to see whether their related diseases or factors tend to concentrate on certain diseases. Among the entries, 131 entries indicate an indirect experiment in validating PS; 50 entries include direct PS-validating experiments while only 4 consists of clinical investigations. (**Figure 1**) These results show that most researches concerning PS still focus on the phenomenon that proteins or other biomolecules condense as droplets. As shown in **Figure 2A**, the majority of PS factors collected belongs to Homo Sapiens, almost doubling the number of PS factors in Mus Musculus. Proteins still took up a large portion of all PS entries, while PS of RNAs also contribute to diseases. (**Figure 2B**) Some key factors like TAR DNA-binding protein 43 (TDP-43) and fused in sarcoma (FUS) appeared in more than 10 entries, indicating the significance PS plays in these key factors concerning neurodegenerative disease. We found that a wide variety of diseases are associated with PS, most of which are nervous system diseases and cancer. (**Figure 2C**) To find out the specific disease that PS has the most effects on, we have also calculated the distribution of PS factors in nervous system diseases and cancer, two of the largest categories where PS play a big part in. We found that among the 80 “nervous system disease” entries, 27 has relations to one rare neurodegenerative disease named amyotrophic lateral sclerosis (ALS), which is even greater than the number of related entries of a more general “neurodegenerative diseases” category (**Table 1**). On the cancer side, diseases distribute more evenly than neurodegenerative disease sector. Lung non-small cell carcinoma has been the leading disease, with 8 entries, while the most entries on the cancer side belongs to “cancer general process” (11 entries) (**Table 2**). These indicates that most cancer researchers tend to investigate the general mechanisms PS plays in cancer. Besides, there are also a number of other diseases that PS has an influence on. For instance, there are 6 entries related to COVID-19, 5 entries related to autosomal dominant polycystic kidney disease, 2 entries related to Noonan syndrome, a genetic disorder with heterogeneous phenotypic manifestations, etc. These results indicated that PS-related factors are widespread among many disorders and PS is a significant mechanism that should not be neglected in investigating these diseases.

**Figure 2.**
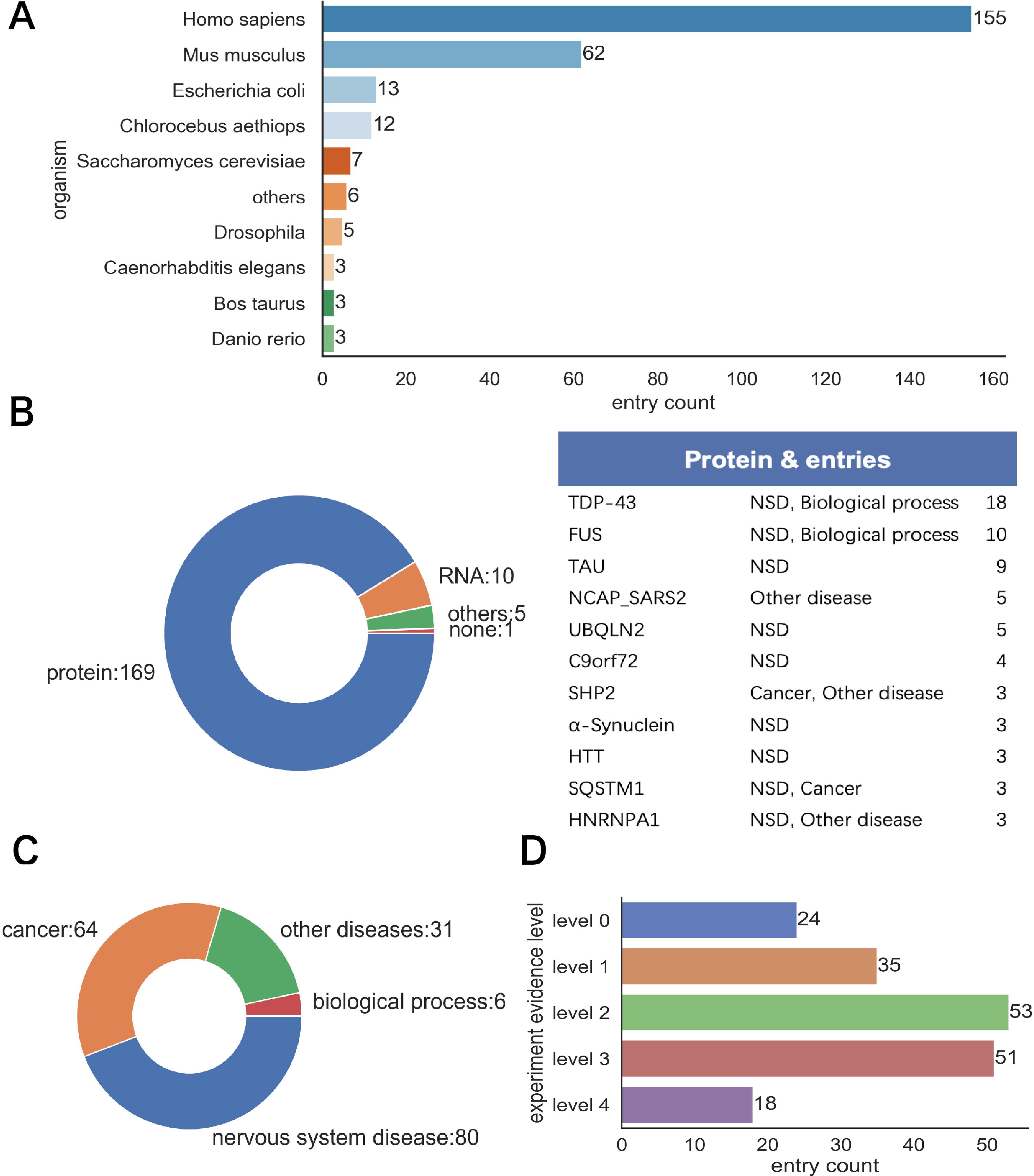
Statistical analysis of entries in PhaSeDis. **A.** Distribution of the organisms of the entries. **B.** Distributions of the factor types of the entries (Left). A list of factors in PhaSeDis with the most appearances in the entries (Right). **C.** Distribution of the diseases associated with each entry. **D.** Entry counts of different levels of phase separation evidence. Level 0 to 4 stands for the number of valid *in vivo* experiments presented in the corresponding literature of each entry. NSD, nervous system disease.

**Table 1.**
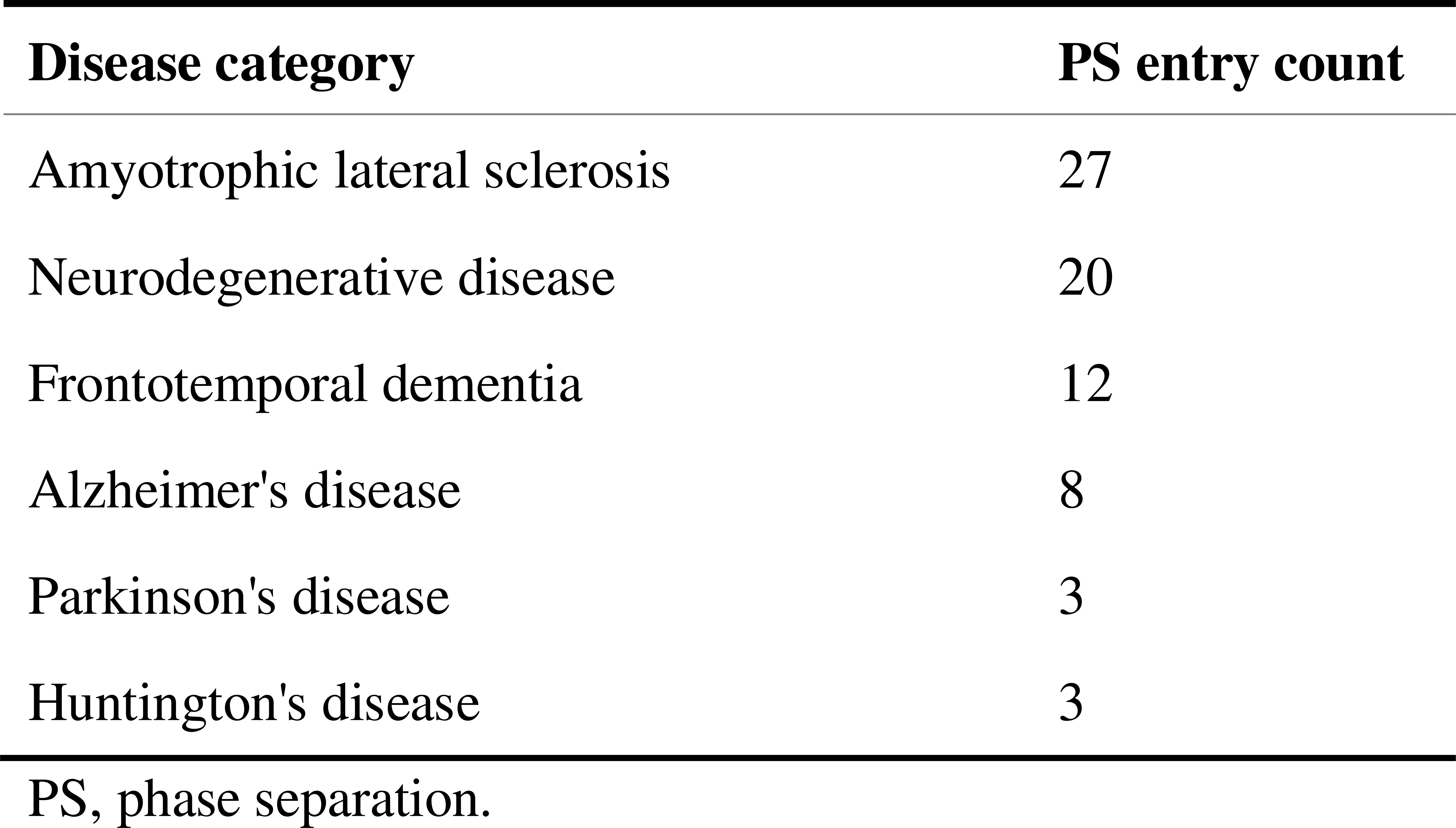
Number of phase separation entries related to nervous system diseases.

**Table 2.**
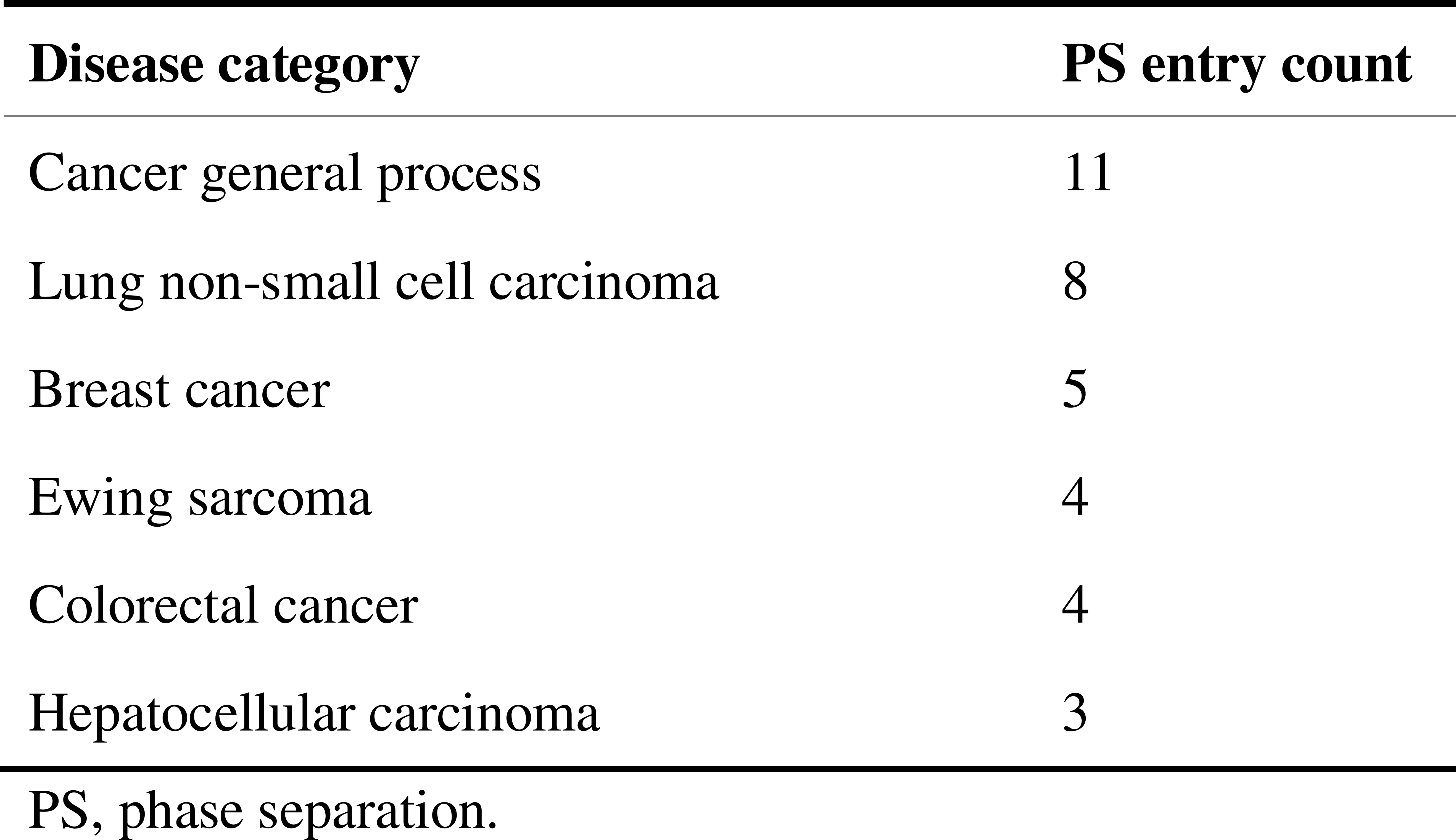
Number of phase separation entries related to cancer.

One of the most concerning problems in the PS field is the verification of PS *in vivo*. As a novel concept in biological context, there is no “golden standard” in determining whether a membraneless organelle or granule forms via PS [12]. Therefore, we supplemented another novel property for each PS entry entitled “*in vivo* PS evidence”. We listed 4 of the most renowned experiments that can help to determine whether the protein can phase separate *in vivo*: 1) fluorescence recovery after photobleaching (FRAP). This experiment labels PS factors with fluorescence and photo-bleach a small spot of area within the condensate with high-energy lasers and record the fluorescence recovery time. FRAP allows for the comparison of viscosity between phase separating droplets. 2) 1,6-hexanediol (1,6-hexa) treatment. 1,6-hexa is one of the most used small molecules that can dissolve PS droplets. Many researches would treat droplets with 1,6-hexa to confirm their LLPS nature [14]. 3) Spherical morphology. The nature of LLPS indicates that the droplet tends to form a circular shape due to surface tension, since no membranes are bound to the droplet. 4) Fusion & fission phenomenon. When two droplets share similar viscosity and composition, they tend to fuse when their boundaries touch. A phase separation droplet can also divide into multiple small droplets. Considering the wide acceptance of these experiments when determining phase separation, we annotated the ‘phase separation’ entries with the existence of said *in vivo* experiment evidences in the corresponding references. The sub-columns are annotated as “Yes” if the corresponding experiment was performed *in vivo* in the reference article mentioned in the entry, and “No” vice versa. Another sub-column named “PS role” indicated the possible role of the corresponding factor in forming the phase separation condensate, which could be “scaffold” if the molecule is the main condensing or recruiting factor, “client” if the molecule is being recruited or transported into the droplet, or “self” if the droplet was condensed by the molecule solely. After annotation we summed up the number of experiments each entry have and named it as the “level of evidence” for the corresponding entry. Each entry ranges from “level 0” (having done none of the aforementioned validation experiments) to “level 4” (having done all validation experiments). As shown in **Figure 2D**, we can see that the majority of entries have a level 2 or level 3 of confidence. Only 18 entries have done all 4 experiments *in vivo* and 24 entries didn’t perform any *in vivo* PS property validations. These results indicate the possible levels of confidence one can expect from a phase separation publication.

### Curation of related small molecules concerning phase separation

Previous researches have shown that small molecules can serve as drugs targeting disease-related proteins [28,29]. In PhaSeDis, we integrated these individual evidences and constructed a “Related small molecules” section for each entry. For each entry, we checked their corresponding reference publication in search of small molecules interacting with or disrupting PS and listed the description in “Interference note” of the “Related small molecules” section. We manually curated how small molecules influence the pathological processes from each reference publications and annotated them in the entry. Among all entries in the database, 28 entries include evidences where small molecules can regulate the pathological processes, and 25 of them are PS entries (**Figure 3**). This information provides low-throughput evidences of small molecule interference to PS *in vivo*.

**Figure 3.**
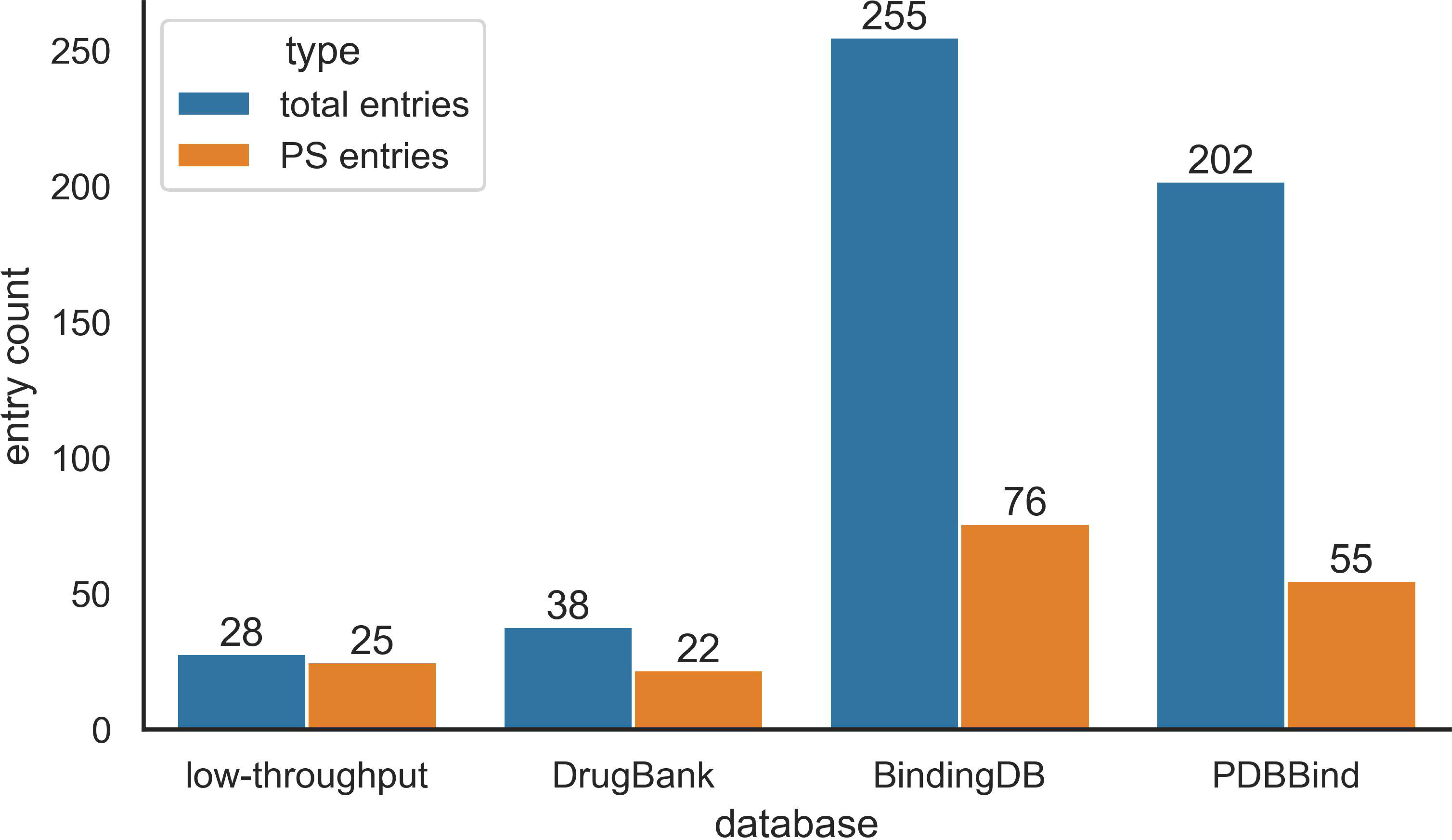
Distribution of related small molecule data in PhaSeDis. Distribution of low-throughput evidences from corresponding literature were illustrated in “low-throughput” column. Counts of high-throughput screening results searched against DrugBank, BindingDB, and PDBBind were illustrated in the corresponding columns.

Besides low-throughput experimental evidences from literature, many databases with information of small molecule–protein interaction data including BindingDB, PDBBind, and DrugBank would also include a wide range of small molecules and their targets [30−33]. Nevertheless, none of these databases would illustrate the significance of PS when it comes to small molecule targets. Therefore, based on our previous PS protein–disease relations curated in PhaSeDis, we searched all PS proteins in our database against these databases. BindingDB provides large amount of protein–small molecule interactions with experimental data. For BindingDB, we searched the UniProt ID of all PS proteins in PhaSeDis and acquired a list of interacting small molecules for each protein. Since most of the molecules do not have a unified common name, we used their PubChem CID as the unique representation of each molecule and annotated them to the corresponding entry as “Molecules from Binding database”. We also searched our PS factor entries against PDBBind, a database with comprehensive information about protein–ligand and other interactions. Besides binding affinity data, PDBBind presents information about the structure details of protein–ligand binding, which increases the liability of protein–ligand interactions. The small molecules derived from PDBBind were annotated to the corresponding entry as “Molecules from PDBBind”. Although having higher tendency to interact with PS factors, molecules found in BindingDB and PDBBind does not necessarily qualify as a potential drug since they do not provide information concerning diseases. Therefore, we also searched all PS proteins, both in Gene ID and UniProt ID, in PhaSeDis against DrugBank, where most small molecules have greater probabilities to qualify as drugs for certain diseases. The interacting drugs related to each protein were annotated in the entry as “Drugs from DrugBank”. DrugBank serves as a database for drugs, indicating that small molecules from DrugBank targeting corresponding PS factors can affect diseases with pre-clinical or clinical evidences. In total, there are 255, 202, and 38 entries whose factors have relations with small molecules in BindingDB, PDBBind, and DrugBank respectively. For verified PS entries, the number of entries with small molecule relations are 76, 55, and 22 for BindingDB, PDBBind, and DrugBank respectively (**Figure 3**). These small molecules are listed in separate columns of the “Related small molecules” section and constitutes as the high-throughput evidences of possible PS-related small molecules.

### Database web interface

The freely available and fully functional website of PhaSeDis (http://mlodis.phasep.pro) has been greatly improved in order to handle the updated information. The website contains six sections, namely ‘Home’, ‘Browse’, ‘MLOs’, ‘Diseases’, ‘About’, and ‘Download’ (**Figure 4A**). Users can search entries of interest on homepage by MLO, factor name, factor gene ID, disease as well as organism or browse all entries on ‘Browse’ page. Users can also choose from all possible options from “Evidence” or “Class” types to browse one specific type of entries (**Figure 4B**). The query results are presented in a single table containing MLOs, factors, diseases, and entry groups (**Figure 4C**). ‘Diseases’ page provides a thorough list of all LLPS/MLO-related diseases illustrated in the database. Clicking on each disease name will bring viewers to all related entries in the database. (**Figure 4D**) ‘MLO’ page includes a straightforward scheme, providing a user-friendly graphical navigation enabling users to browse all entries related to an interested MLO by simply clicking on the MLO (**Figure 4E**). There are also ‘MLO detail’ page providing basic introduction to every MLO (**Figure 4F**). Users can click on ‘LLPS’ on the ‘MLOs’ page to see all LLPS–disease entries with confirmed LLPS proteins.

**Figure 4.**
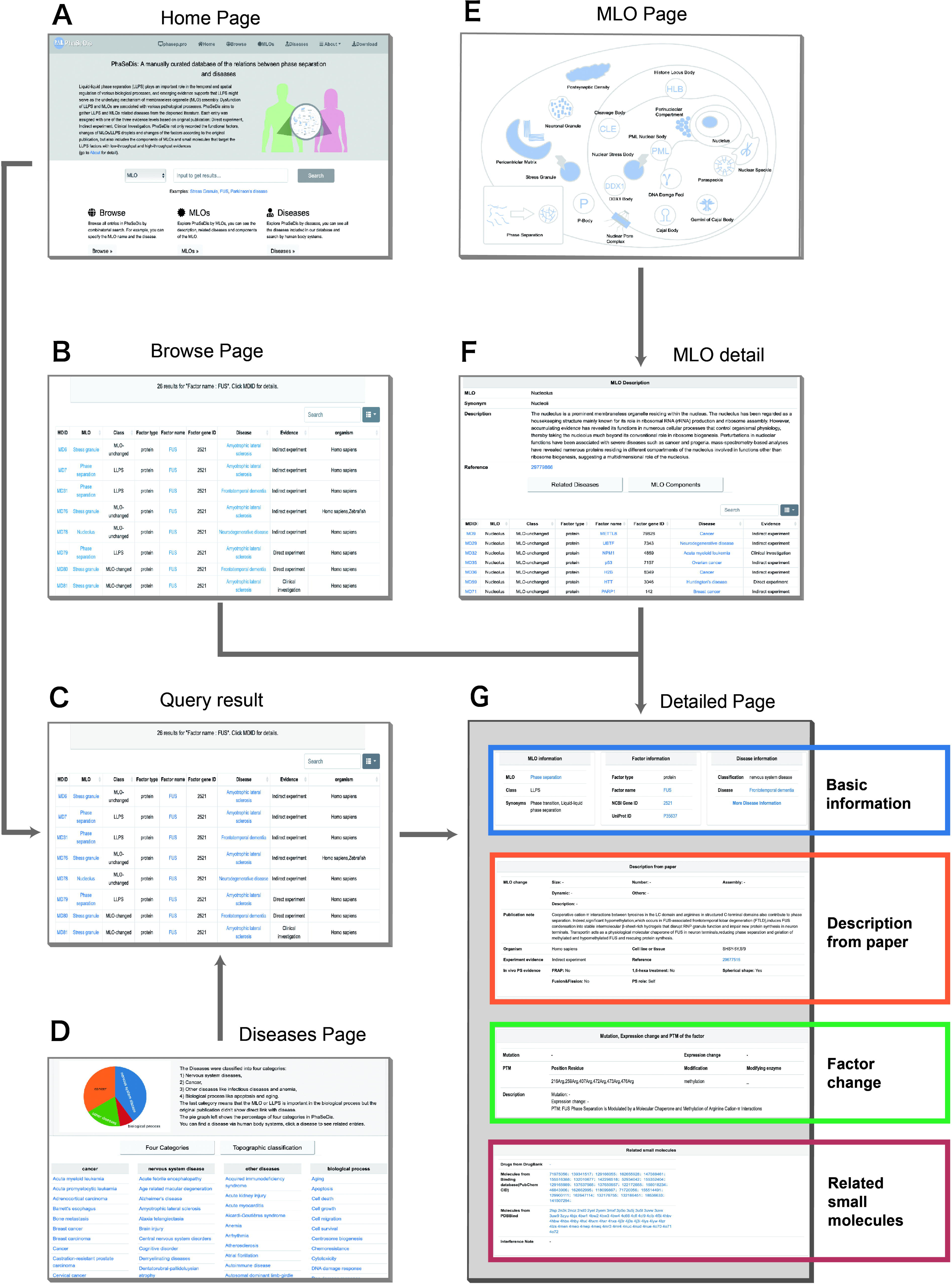
The web interface of PhaSeDis. **A.** Home page. Users can search by MLO, factor name, factor gene ID or disease. **B.** Browse page. Users can browse entries of interest in a simple table form. **C.** MLO page. Users can navigate the MLOs through a graphical scheme. **D.** Disease page. Users can see all the diseases collected in PhaSeDis in classified tables. **E.** Detailed page. This page shows all the information contained within an entry. MLO, membraneless organelle.

For every entry we assigned a unique in-built ID named MDID, which you can see in ‘Browse’ page, ‘Query result’ page or ‘MLO detail’ page. Clicking on any MDID would bring viewers to the detail page of the entry (**Figure 4G**). The detail page of each entry consists of 4 parts: 1) Basic information, which contains the information of MLO, factor, and disease. 2) Description from paper, which encompasses the changes in MLO or LLPS bodies in terms of size, number, assembly conditions, dynamic, and others, including a “Publication note” with curation of how condensation might participate in pathological processes. This section also includes the PubMed ID of the original article, as well as the organism, cell/tissue type, experimental evidence and a brief introduction of the results on the paper. For LLPS entries, there’s also annotations of whether *in vivo* PS evidence exists, as well as the role the corresponding factor play in PS (Self/Scaffold/Client). 3) Mutation, expression change, and post-translational modification (PTM), including all mutations and changes occurring in the original article and their impact. 4) Related small molecules. This section illustrates both low-throughput and high-throughput evidences of targeting or interacting small molecules. The low-throughput evidences regarding the impact of related small molecules on cellular or organismal pathological conditions are collected from literature sources and detailed in the “Interference notes” section. These indicate the potential changes observed when the small molecules are introduced. Conversely, small molecules supported by high-throughput evidences are systematically linked to their corresponding databases.

User guide and data summary are detailed in ‘About’ page. In ‘Download’ page, we added a “PhaSeDis data” item, with the description “This table contains all entries in PhaSeDis, including information about related small molecules”, so that users can download all data in PhaSeDis freely for further analysis.

## Discussion

Phase separation can help explain the concentration and sequestration of certain proteins or nucleic acids, which can accelerate or inhibit certain biological functions and form membraneless organelles. Dysregulation of phase separation is an important pathological mechanism, particularly in nervous system diseases and cancer. However, due to a lack of “golden standard”, researchers do not have a consensus of the detrimental experimental observation and illustrate the PS properties of proteins with different sets of methodologies. In this work, we updated the phase separation entries in PhaSeDis to include the precise experimental evidence of the factor in the original publications, allowing a more comprehensive view on the reliability of each entry. We have exclusively included *in vivo* PS evidences because such data are more reflective of actual physiological conditions within a biological context compared to *in vitro* experiments.

Besides, PS properties of biomolecules poses new challenges to drug discovery, since the small molecule might have to partition into or dissolve the condensate to have an impact on the diseases [8]. Many literatures have issued the influence small molecule treatment has on diseased cells or animal models [16,20,34]. In PhaSeDis, we collected these low-throughput evidences illustrated by the literature, as well as high-throughput evidences from protein–small molecule interaction databases including BindingDB, PDBBind, and DrugBank. These databases provide possible small molecule interaction data from different aspects: BindingDB focuses on the binding nature of small molecules with PS factors that are related to diseases; PDBBind includes a variety of small molecules/ligands binding to proteins at specific regions, which can be displayed as 3D structures and stored in PDB files; DrugBank provides a list of approved small molecules that are more likely to deal with the disease from the respective entry. We do not intend to imply that these small molecules necessarily bind to or interfere with PS factors through any specific mechanism, nor do we suggest that these molecules themselves undergo phase separation. In fact, research suggests that certain small molecules might not operate in the conventional manner of drugs that bind to target pockets; rather, they could potentially exert their disease-modifying effects through noncanonical modes of action [35]. BindingDB, PDBBind, and DrugBank themselves are not databases on PS, thus providing no direct links between PS and small molecules. Nevertheless, these small molecules derived from protein–small molecule interaction databases have a higher probability of interfering with PS since they target these key PS factors. Besides small molecule–protein interaction databases, there are also databases that introduces the relations between small molecule perturbance and gene expression changes in a variety of diseases, e.g. cMAP [36]. Nevertheless, the relationship between gene expression changes and protein binding is complex and not straightforward to establish. Therefore, as a database specifically dedicated to the relationship between PS (proteins/RNA) and diseases, PhaSeDis does not incorporate data from cMAP.

In recent literature, there are also other databases that focus on the relationship between PS and diseases, including DisPhaseDB [37]. Although PhaSeDis and DisPhaseDB both include information about PS protein and disease relations, there are some differences between the two databases. DisPhaseDB collects proteins with LLPS evidence from databases and collects related diseases based on the mutations in ClinVar, with a focus on “variants”. PhaSeDis collects PS factors and their relations to diseases in existing literatures, as well as potentially interfering small molecules from literature and databases. As shown in **Figure S1**, there are 231 proteins overlapping between the two databases, however, PhaSeDis contributes an additional 117 unique proteins that are not covered in DisPhaseDB, taking up roughly one third of all PhaSeDis entries (**Figure S1**). PhaSeDis is also a great supplement to existing PS factor databases, e.g., PhaSePro [24]. As shown in **Figure S2**, PhaSeDis and PhaSePro has only 25 proteins in common, with 323 unique in PhaSeDis. PhaSeDis incorporates more recent research findings and specifically dedicated to diseases where PS factors play a role and how small molecules can intervene in these processes, taking a distinct approach compared to other PS databases that primarily concentrate on cataloguing LLPS proteins.

Currently, summarizing mechanisms of diseases concerning PS systematically, as well as mechanisms of small molecules targeting PS, are still challenging and does not rely on any single properties [38,39]. Researchers dedicated to different diseases have their own hypothesis in how PS affects disease progression and alleviation [10,40]. For PS-intervening small molecules, some researchers have put forward a few classifications of mechanisms [41], and our data can partially fit in: 1) Some molecules work as dissolvers, like in MD787, where phospholipase D inhibitor 1-Butanol or 2-Butanol decrease large tumor suppressor kinase 1 (LATS1) puncta numbers [15]; 2) Some molecules work as inducers, like in MD792, where sulforaphane (SFN) treatment doubles the number of super-enhancers to increase nuclear factor erythroid 2-related factor 2 (NRF2) activity and improve kidney function for autosomal dominant polycystic kidney disease (ADPKD) patients [16]; 3) Some molecules work as localizers, like in MD838, where 5-fluorouridine and ebselen targets superoxide dismutase 1 (SOD1) W32S and C111S sites to prevent oxidation-induced aggregation [17]; 4) Some molecules work as morphers, like in MD851, where lipid incorporation into α-synuclein droplets delays their maturation process, thereby rendering the droplets more resistant to aging [18]. However, other types of mechanisms have also been proposed [9]. For instance, it could also stem from the conformational changes induced by drug binding, which effectively interfere with and hinder the progression of pathological conditions [42], as in MD814–MD816, where Src homology region 2 domain-containing phosphatase-2 (SHP2) allosteric inhibitor attenuates LLPS of SHP2 mutants [29]. As the currently available evidence does not permit us to draw definitive conclusions about the specific nature of drug–PS proteins–disease interactions, PhaSeDis aims to provide detailed introduction to every entry concerning how PS factors lead to pathological conditions through phase separation, and how small molecules intervene. As PhaSeDis accumulates more data, we hope that it will facilitate deeper understanding of the *in vivo* roles of PS phenomena, their associations with diseases, and the identification of potential therapeutic agents. For now, the current version of PhaSeDis serves as a valuable starting point for initiating a multitude of research inquiries into drug–PS protein–disease interactions.

There are also a few limitations on this work: firstly, the number of PS factors are limited, which makes it difficult to summarize and categorize the features of the PS-related pathological mechanisms. Secondly, the protein–small molecule databases from which we source our small molecules do not specifically emphasize PS aspects. Therefore, researchers need to further investigate and validate the mechanisms by which these small molecules potentially influence PS.

Taken together, PhaSeDis has grown into a comprehensive database for PS factor–disease relations, as well as the first database of its kind that integrates small molecule interaction in the database. These invaluable resources will greatly assist researchers who have a keen interest in the realm of PS, empowering them not only to rigorously substantiate the intricate partitioning mechanisms of currently available medications but also to embark on pioneering quests for novel therapeutic agents capable of effectively disrupting PS. Such endeavors hold immense promise for advancing our understanding of PS, ultimately contributing to the development of innovative treatment strategies.

## Data availability

PhaSeDis is publicly available at http://mlodis.phasep.pro.

## CRediT author statement

**Taoyu Chen:** Conceptualization, Methodology, Formal analysis, Investigation, Data curation, Writing – Original Draft. **Guoguo Tang:** Formal analysis, Investigation, Data curation, Writing – Review & Editing, Visualization. **Tianhao Li:** Methodology, Software, Investigation, Formal analysis, Visualization. **Zhining Yanghong**: Software, Formal analysis, Visualization. **Chao Hou:** Software. **Zezhou Du:** Data curation. **Liwei Ma**: Writing – Review & Editing. **Tingting Li:** Conceptualization, Methodology, Writing – Review & Editing, Supervision, Project administration, Funding acquisition.

## Acknowledgements

This study was supported by the National Natural Science Foundation of China (Grant Nos T2325003, 32070666, 32100507, 62250005, 31000609), the National Key R&D Program of China (2023YFF1204703, 2021YFF1200900), and the Beijing Natural Science Foundation (Z230014), China.

## Supplementary material

**Figure S1 Venn diagram of the distinction between the proteins contained within DisPhaseDB and PhaSeDis databases.**

There are 231 proteins overlapping between the two databases, while 117 proteins are unique to PhaSeDis and 5510 proteins are unique to DisPhaseDB.

**Figure S2 Venn diagram of the distinction between the proteins contained within PhaSePro and PhaSeDis databases.**

There are 25 proteins overlapping between the two databases, while 323 proteins are unique to PhaSeDis and 96 proteins are unique to PhaSePro.

