## Supplementary figures and images for "PhaSeDis: A Manually Curated Database of Phase Separation–Disease Associations and Corresponding Small Molecules"

### Figure S1

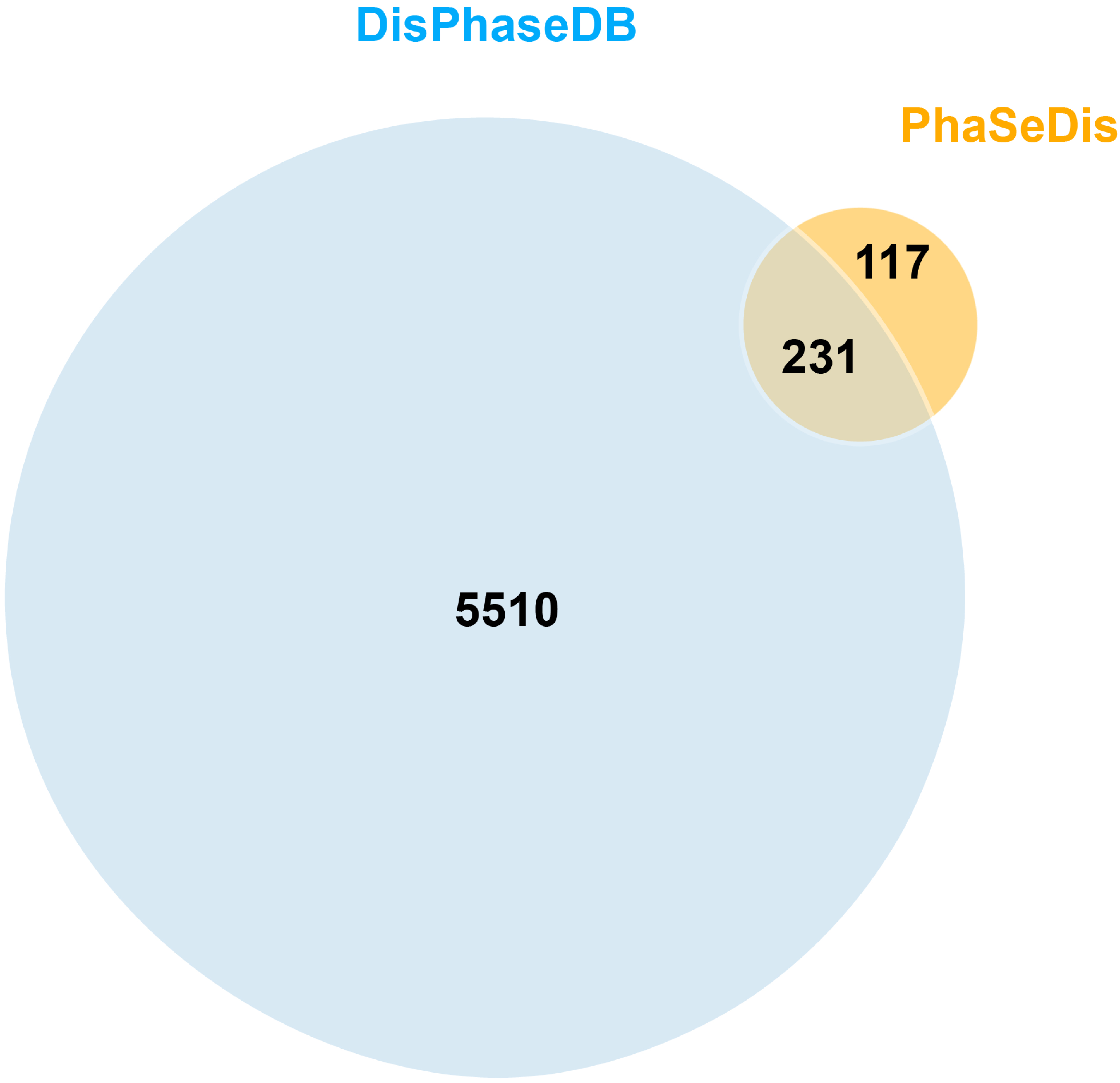

### Figure S2

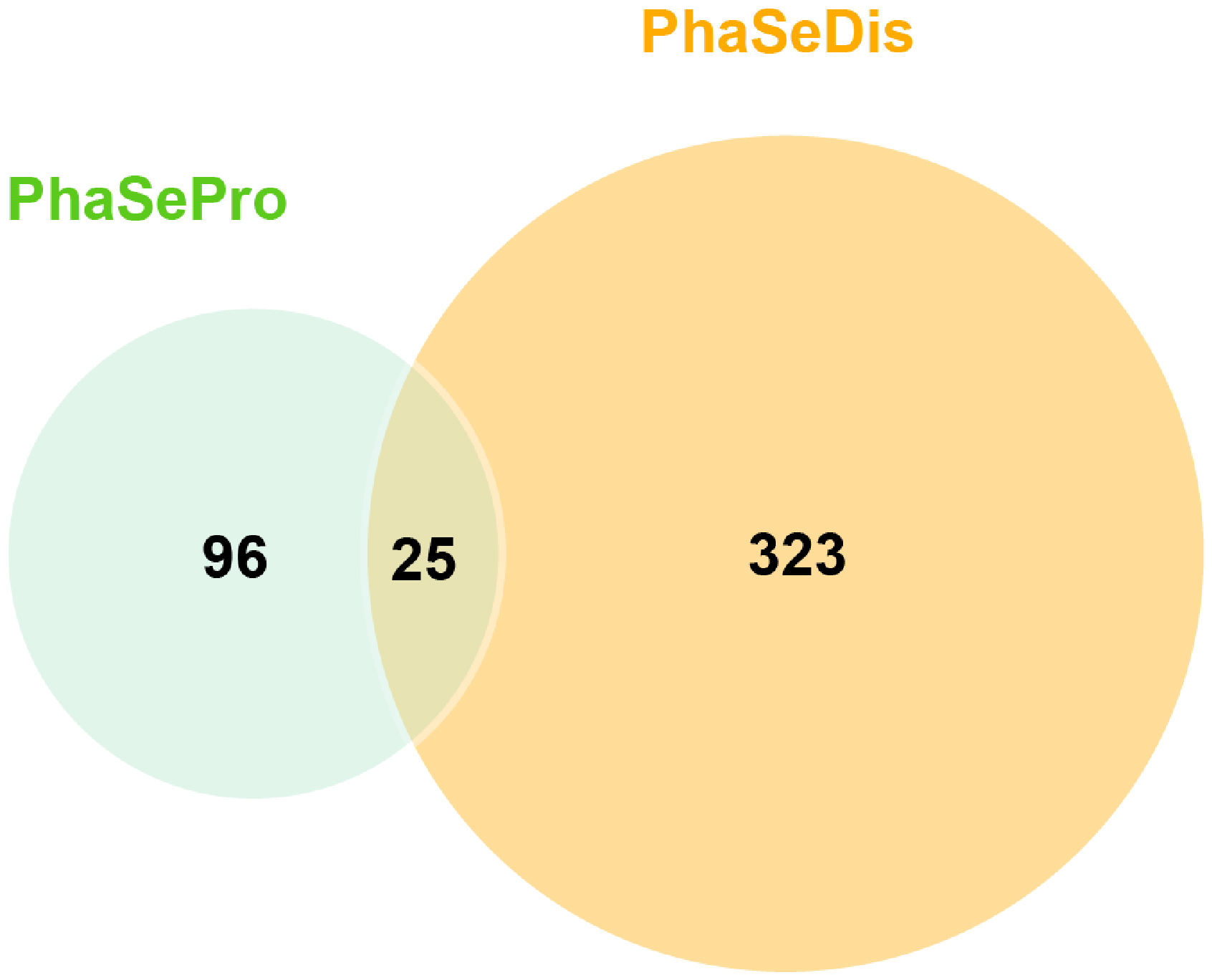
